# A Virus-like particle-based bivalent PCSK9 vaccine lowers LDL-cholesterol levels in Non-Human Primates

**DOI:** 10.1101/2023.05.15.540560

**Authors:** Alexandra Fowler, Koen K. A. Van Rompay, Maureen Sampson, Javier Leo, Jennifer K. Watanabe, Jodie L. Usachenko, Ramya Immareddy, Debbie M. Lovato, John T. Schiller, Alan T. Remaley, Bryce Chackerian

## Abstract

Elevated low-density lipoprotein cholesterol (LDL-C) is an important risk factor in the development of atherosclerotic cardiovascular disease (ASCVD). Inhibitors of proprotein convertase subtilisin/kexin type 9 (PCSK9), a negative regulator of LDL-C metabolism, have emerged as promising approaches for reducing elevated LDL-C levels. Here, we evaluated the cholesterol lowering efficacy of virus-like particle (VLP) based vaccines that target epitopes found within the LDL receptor (LDL-R) binding domain of PCSK9. In both mice and non-human primates, a bivalent VLP vaccine targeting two distinct epitopes on PCSK9 elicited strong and durable antibody responses and lowered cholesterol levels. In macaques, a VLP vaccine targeting a single PCSK9 epitope was only effective at lowering LDL-C levels in combination with statins, whereas immunization with the bivalent vaccine lowered LDL-C without requiring statin co-administration. These data highlight the efficacy of an alternative, vaccine-based approach for lowering LDL-C.

## Introduction

Cardiovascular disease (CVD) is the leading cause of global mortality, responsible for approximately 19 million deaths in 2020^1^. A major risk factor for atherosclerotic cardiovascular disease (ASCVD) is elevated plasma levels of low-density lipoprotein cholesterol (LDL-C)^2, 3^. Correspondingly, reducing levels of circulating LDL-C can lower the risk of ASCVD^4, 5^. Statins are the most commonly prescribed medication for lowering LDL-C levels. Statins, which inhibit 3-hydroxy-3-methyl-glutaryl coenzyme A (HMG-CoA) reductase, an enzyme in the cholesterol biosynthetic pathway, can decrease both LDL-C levels and the risk of cardiovascular events^6^, but their effectiveness in lowering LDL-C varies amongst individuals. As many as 20% of patients are hypo-responsive to statin use^7, 8^. Although statins are generally well-tolerated, their use can be associated with serious adverse effects, including myopathy and liver toxicity^9^. These limitations have prompted the development of non-statin lipid-lowering therapies that target other pathways involved in LDL-C metabolism^10^.

LDL-C is removed from circulation by the low-density lipoprotein receptor (LDL-R), which is most abundantly expressed in the liver. Once LDL-C interacts with LDL-R the complex is endocytosed. The low pH of the endosome causes the complex to dissociate, allowing LDL-C to be metabolized by the cell and LDL-R to be recycled back to the plasma membrane where it can continue to remove circulating LDL-C. Proprotein convertase subtilisin/kexin type 9 (PCSK9) is a serum-associated secretory protein that directly inhibits the recycling of LDL-R. PCSK9 binds to the extracellular domain of LDL-R and this strong interaction mediates its lysosomal degradation upon endocytosis^11^. Thus, by inhibiting LDL-R recycling, PCSK9 mediates higher levels of circulating LDL-C. Genetic mutations that affect PCSK9 activity can have profound effects on LDL-C levels. Patients with gain of function (GOF) mutations that increase the affinity of PCSK9 for LDL-R can develop autosomal dominant hypercholesterolemia (ADH); these patients exhibit high levels of LDL-C and early onset ASCVD^12^.

Conversely, patients with loss of function (LOF) mutations in PCSK9 exhibit low LDL-C levels and a decreased risk of cardiovascular events^13, 14^. Importantly, human LOF mutations are not associated with any adverse consequences. Thus, PCSK9 has become a major therapeutic target for lowering circulating LDL-C and preventing ASCVD.

Several different strategies for inhibiting PCSK9 activity have been developed, including monoclonal antibodies (mAbs), small interfering RNA (siRNA), macrocyclic peptides, base editing, and vaccines^15^. Currently, there are three FDA-approved PCSK9 therapies that are effective at reducing LDL-C levels in combination with statins. Evolocumab (Repatha^®^) and alirocumab (Praluent^®^) are anti-PCSK9 mAbs that lower LDL-C levels by as much as 50-60%^16–18^. Inclisiran (Laqvio^®^) consists of liposomes encapsidating an siRNA cargo that prevents production of PCSK9 in the liver. Inclisiran is similarly effective to mAbs, reducing LDL-C by 40-60%^19^, but unlike mAbs, which need to be administered every 2-4 weeks, Inclisiran has long-lasting effects and only needs to be administered twice a year. However, both mAb- and siRNA-based PCSK9 inhibitors are expensive; alirocumab and evolocumab cost approximately US$6000 annually and inclisirin costs US$3250 per dose^20^, leading some to question the cost effectiveness of these drugs^21–23^. Because of their expense, anti-PCSK9 therapies are mostly used as secondary prevention in patients who do not achieve sufficiently low LDL-C levels using statins alone^24^.

Vaccines are another promising approach for modulating PCSK9 activity. Vaccines have several potential advantages over other therapeutic approaches; they are relatively inexpensive to produce, which could reduce patient costs, and they will likely require fewer doses, potentially increasing patient compliance. Although the immunological mechanisms of self-tolerance normally restrict the ability to induce antibody responses against self-antigens such as PCSK9, these mechanisms can be efficiently overcome by displaying self-antigens at high density on the surface of nanoparticle-based vaccine platforms, such as virus-like particles (VLPs)^25–27^. VLP-based vaccines targeting self-antigens have been evaluated in human clinical trials, and this approach has been shown to be safe and able to induce high titer antibody responses^28, 29^. In previous work, we engineered VLP-based vaccines that displayed different linear peptides from human PCSK9 that were predicted to interact with LDL-R^30^. We identified several vaccine candidates that induced high titer anti-PCSK9 antibody responses and reduced total cholesterol levels in immunized mice. In this study, we have engineered and combined VLP-based vaccines that display two species-specific linear peptides from PCSK9 to test their efficacy to induce anti-PCSK9 antibody responses and reduce cholesterol levels in multiple animal models. Our results support the use of a bivalent VLP-based vaccine targeting two epitopes of PCSK9 to effectively lower LDL-C levels without requiring co-administration of statins.

## Results

### Evaluating the immunogenicity and efficacy of single and bivalent PCSK9 vaccines in the LDLR^+/-^ mouse model

We previously showed that VLP-based vaccines targeting linear epitopes from human PCSK9 (hPCSK9) elicit high-titer anti-PCSK9 antibodies and lower cholesterol levels in mice^30^. Although the amino acid sequences of hPCSK9 and mouse PCSK9 (mPCSK9) are highly conserved, there are several differences within the epitopes we targeted that potentially could affect the anti-PCSK9 activity of induced antibodies (Fig. 1). In addition, our original study did not assess the effectiveness of bivalent (combination) vaccines targeting multiple PCSK9 epitopes. To address these issues, we produced VLPs displaying mPCSK9 peptides that correspond to the hPCSK9 epitopes that, when displayed on VLPs, most potently decreased cholesterol levels in Balb/c mice^30^. These epitopes (amino acids 153-163 and 207-223) are located on the face of PCSK9 that is involved in LDL-R binding (Fig. 1). Peptides representing mPCSK9_153-163_ and mPCSK9_207-223_ were synthesized and then conjugated onto Qß bacteriophage VLPs using a bifunctional chemical cross-linker. Each peptide was displayed on VLPs at high valency, ∼360 peptides per VLP (data not shown).

**Fig 1.**
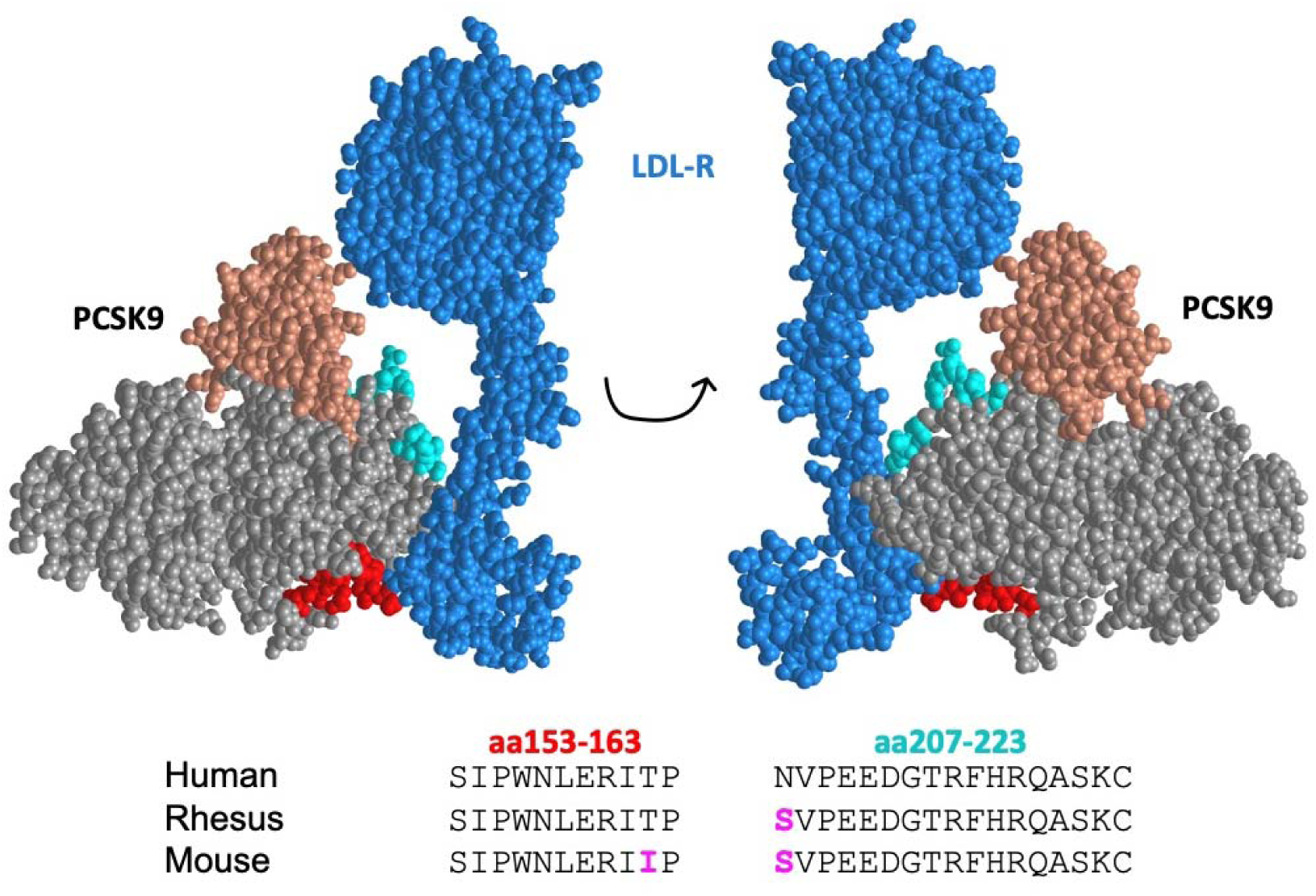
The structure of human PCSK9 in complex with LDL-R. LDL-R is shown in blue and the prodomain and catalytic domain of PCSK9 are shown in salmon and grey, respectively. Targeted PCSK9 epitopes are shown in red (amino acids 153-163) and cyan (amino acids 207-223). The amino acid sequences of these epitopes from human, rhesus macaque, and mouse PCSK9 are shown; residues that differ from the human sequence are shown in magenta. Structures are generated using the RCSB Protein Data Bank, structure ID: 3M0C.

To evaluate the immunogenicity and cholesterol lowering activity of mPCSK9-targeted vaccines, we utilized mice heterozygous for the *Ldlr^tm1Her^*mutation which express low levels of functional LDL-R and have elevated serum cholesterol^31^. We had evaluated vaccines targeting the homologous epitopes from hPCSK9 previously^30^; to confirm that VLPs displaying mPCSK9 epitopes were similarly able to elicit anti-PCSK9 antibody responses, a small group of *LDLR^+/-^* mice (n=6) were immunized three times with 5µg doses of mPCSK9_153-163_ VLPs, mPCSK9_207-223_ VLPs, or, as a negative control, wild-type Qß VLPs. In addition, a larger group of mice (n=18; with a similarly sized group of control mice) were immunized with a bivalent PCSK9 vaccine, which consisted of a mixture of 5µg of mPCSK9_153-163_ VLPs and 5µg of mPCSK9_207-223_ VLPs. The immunogenicity of each vaccine was evaluated by measuring antibody titers to the target PCSK9 peptides and to full-length recombinant mPCSK9. As is shown in Fig. 2, both individual and bivalent VLP-based vaccines elicited high titer IgG antibody responses against mPCSK9_153-163_, mPCSK9_207-223_, and full-length mPCSK9. Although the bivalent vaccine elicited slightly lower anti-peptide antibody titers than the individual vaccines that specifically targeted each epitope (Fig. 2a & 2b), it generated similarly strong anti-mPCSK9 IgG antibody titers (Fig. 2c).

**Fig. 2.**
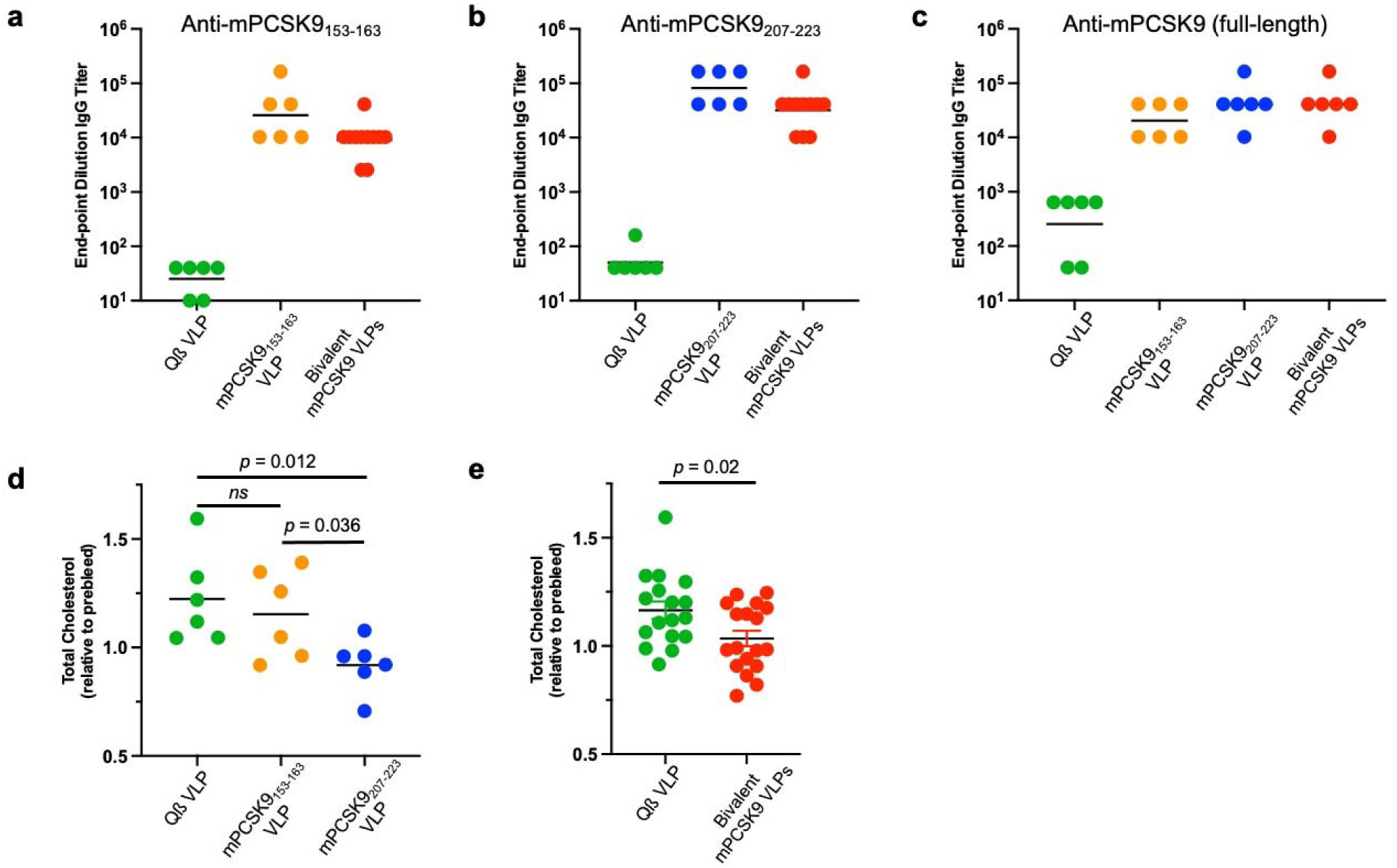
Immunogenicity and cholesterol lowering activity of PCSK9 VLPs in LDLR^+/-^ mice. LDLR^+/-^ mice were immunized three times at three-week intervals with 5µg of wild-type Qß VLPs, mPCSK9_153-163_ VLPs, mPCSK9_207-223_ VLPs, or a bivalent mPCSK9 vaccine consisting of a mixture of 5µg mPCSK9_153-163_ VLPs and 5µg mPCSK9_207-223_ VLPs. Blood plasma were obtained three weeks following the final immunizations and IgG antibody titers against (**a** & **b**) mPCSK9 peptides and (**c**) full-length recombinant mPCSK9 were determined by ELISA. The effects of vaccination on total cholesterol levels were determined by comparing the serum cholesterol level in plasma following the vaccination series with levels in plasma obtained prior to immunization (**d** & **e**). Experimental groups were compared statistically by unpaired two-tailed t test. ns, not significant.

To evaluate the cholesterol-lowering effects of the PCSK9 vaccines in the *LDLR^+/-^* mouse model, we measured the total cholesterol levels of vaccinated and control groups prior to the first immunization and three weeks following the third immunization. As we previously reported in mice immunized with VLPs displaying an analogous hPCSK9 epitope^30^, mice immunized with mPCSK9_207-223_ VLPs had significantly reduced total cholesterol levels (∼25% lower) relative to control mice immunized with Qß VLPs (Fig. 2d). The bivalent vaccine (Fig. 2e) also significantly lowered total cholesterol levels by a similar percentage. Although immunization with mPCSK9_153-163_ VLPs decreased total cholesterol levels relative to controls, this effect was not statistically significant.

### The bivalent PCSK9 vaccine decreases plasma PCSK9 levels in LDLR^+/-^ mice

We and others have shown that anti-PCSK9 antibodies can increase circulating levels of PCSK9 in the serum^30, 32–34^, suggesting that these antibodies may form immune complexes that increase the half-life of serum-associated PCSK9 but interfere with PCSK9 function. We were interested in determining how a vaccine that targets two epitopes might affect plasma PCSK9 levels. As is shown in Fig. 3a, serum PCSK9 levels in mice immunized with VLPs displaying individual mPCSK9 epitopes were either unchanged (mPCSK9_153-163_) or elevated (mPCSK9_207-223_) relative to control mice.

**Fig. 3.**
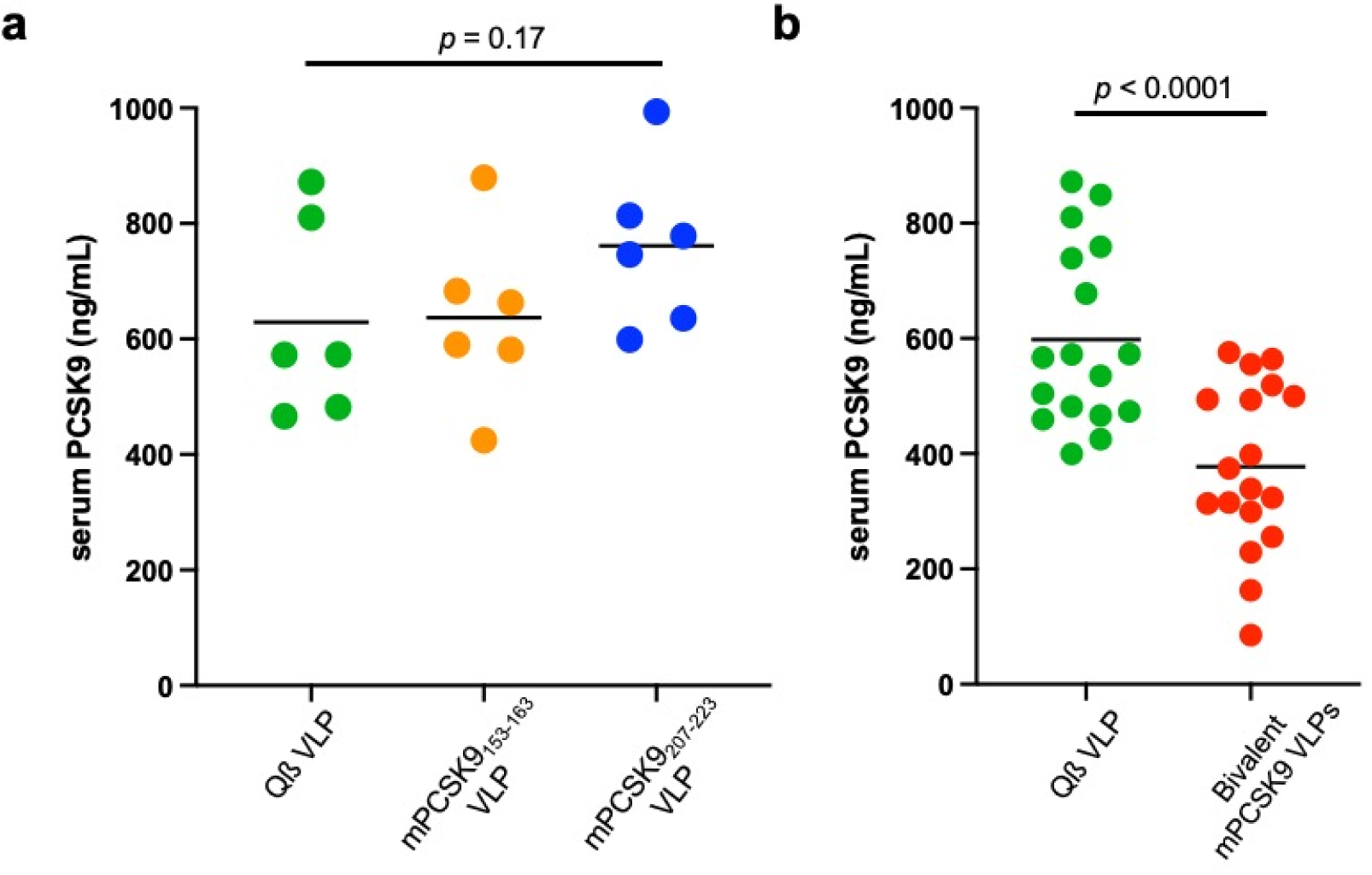
Plasma PCSK9 levels in vaccinated LDLR^+/-^ mice. Mice were immunized three times and plasma were collected two weeks following the final immunization. **a** Plasma PCSK9 levels in mice immunized with individual PCSK9 VLP vaccines or control, wild-type Qß VLPs. **b** Plasma PCSK9 levels in mice immunized with a bivalent PCSK9 VLP vaccine or control, wild-type Qß VLPs. Experimental groups were compared statistically by unpaired two-tailed t test.

However, serum PCSK9 levels were reduced by ∼40% in mice immunized with the bivalent mPCSK9 VLP vaccine (Fig. 3b). Thus, these data confirm Goksøyr’s results and further show that PCSK9 clearance can be mediated by eliciting antibodies that target two distinct epitopes on the protein.

### Immunization with a bivalent PCSK9 VLP vaccine increases liver LDLR expression in LDLR^+/-^ mice

Anti-PCSK9 antibodies are predicted to inhibit PCSK9-mediated LDL-R degradation, resulting in increased LDL-R expression levels in the liver. To evaluate this possibility, livers were harvested from mice immunized with the bivalent mPCSK9 VLP vaccine or control Qß VLPs and LDL-R levels in liver lysates were measured by Western blot analysis. As is shown in Fig. 4, there was a ∼50% increase in LDL-R expression in the livers of mice that were immunized with the bivalent mPCSK9 VLP vaccine, as compared to mice immunized with control Qß VLPs. These data indicate that anti-PCSK9 antibodies functionally inactivate PCSK9.

**Fig. 4.**
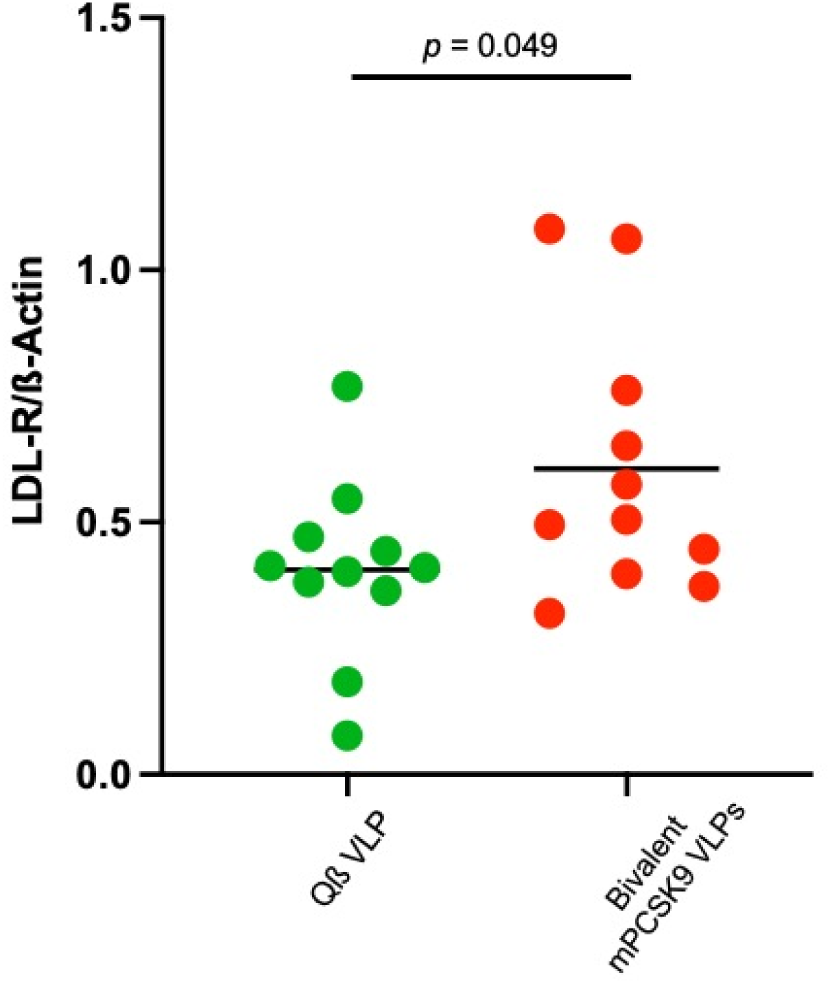
Liver LDL-R expression in immunized mice. LDLR^+/-^ mice (n=11) were immunized three times with a bivalent mPCSK9 VLP vaccine or, as a control, wild-type Qß VLPs. Three to four weeks following the final immunization, mice were euthanized and livers were collected. Liver LDL-R protein expression was measured by Western blot (shown in Fig. S1), quantified using Image J, and then normalized by comparing the expression of LDL-R to β-actin (as a loading control). Each data point represents an individual mouse (n=11) and means for each group are shown using a black line. Experimental groups were compared statistically by unpaired two-tailed t test.

### PCSK9 VLPs elicit long-lived antibody responses

To determine the longevity of the anti-PCSK9 antibody response, we followed a subset of the mice immunized with the bivalent mPCSK9 VLP vaccine for over a year after the initial vaccination. Sera were regularly collected during this period and anti-mPCSK9 IgG titers were measured by ELISA. As is shown in Fig. 5, anti-PCSK9 IgG titers peaked ∼5 weeks following the third immunization. After an initial decline in titers, antibody decay appears to slow substantially, consistent with a pattern of biphasic antibody decay that has been reported for other vaccines^35^. One year after the initial vaccination anti-PCSK9 antibody levels were ∼10-fold lower than peak titers but were still high. Following the initial decline in antibody levels (by week 20), we calculate that the half-life of anti-PCSK9 antibodies induced using the bivalent mPCSK9 VLP vaccine is ∼20 weeks.

**Fig. 5.**
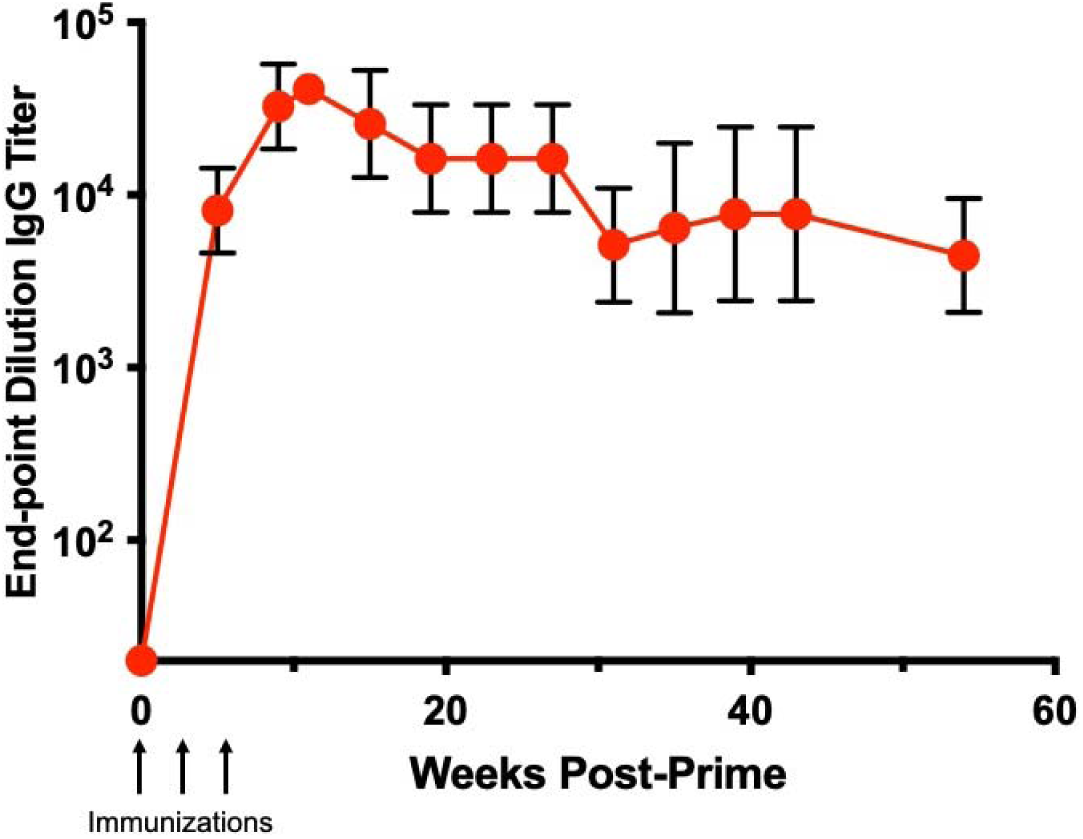
Durability of anti-PCSK9 antibody responses. LDLR^+/-^ mice (n=6; 3 males and 3 females) were immunized three times (at weeks 0, 3, and 6) with a bivalent mPCSK9 VLP vaccine and anti-PCSK9 antibody titers were measured by ELISA for over 1-year post-immunization. Data represents geometric mean end-point dilution titers, error bars show the geometric standard deviation. Note: because of a medical issue (that was unrelated to vaccination) we were unable to obtain sera from one of the mice after week 35.

### Immunization with PCSK9 VLPs elicits strong antibody responses in non-human primates (NHPs)

Non-human primates (NHPs) have a similar lipid profile as humans and have been used as a model for evaluating the efficacy of PCSK9 inhibitors^36, 37^. We assessed the ability of VLP-based PCSK9 vaccines to induce anti-PCSK9 antibody responses and to lower LDL-C in rhesus macaques. 24 healthy rhesus macaques (*Macaca mulatta*) of Indian origin, ranging from 7-12 years old, were identified by the California Regional Primate Research Center. NHPs were pre-screened for total cholesterol and triglyceride levels and were then assigned to three groups (n=8/group) with similar mean lipid levels, weights, and ages (Table S1). After the study was initiated, one of the macaques in the control group (ID: 3-G) developed a medical problem and was withdrawn from the study, resulting in 7 animals in this group. Groups received either (1) a bivalent vaccine consisting of rhesus macaque (rh) PCSK9_207-223_ VLPs mixed with rhPCSK9_153-163_ VLPs (25µg of each), (2) rhPCSK9_207-223_ VLPs alone (50µg), or, as a control group, (3) wild-type Qß VLPs (50µg) at days 0, 28, and 56, as is shown in Fig. 6a. Macaques also received a booster immunization at day 161, approximately four weeks prior to necropsy. Plasma and sera were obtained prior to the first vaccination, after each dose, and then regularly thereafter. The health of immunized macaques was monitored throughout the experiment, and all of the vaccines had a good safety profile, as based on clinical observations, complete blood count (CBC), and standard serum chemistry panels.

**Fig. 6.**
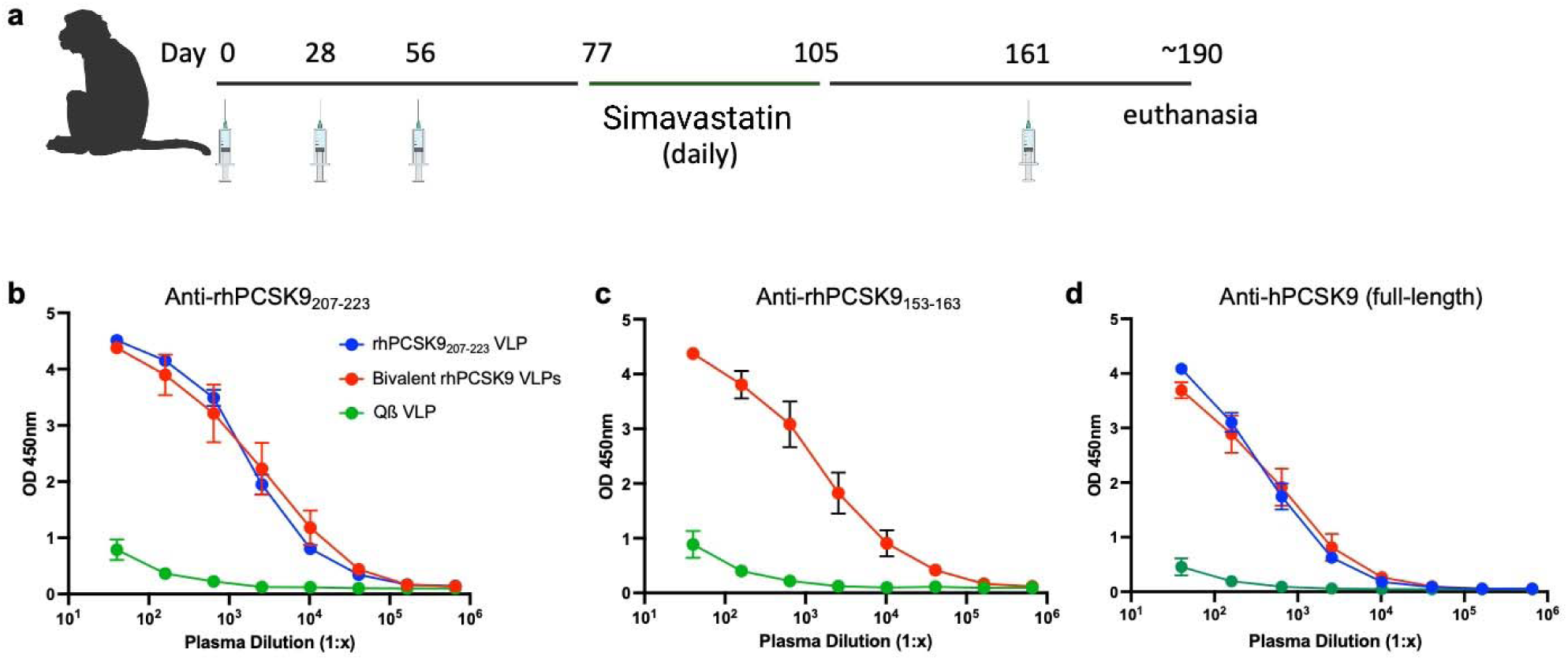
Anti-PCSK9 antibody responses in immunized rhesus macaques. **a** The study timeline. Groups of macaques (n=7-8) were immunized three times (at weeks 0, 4, and 8) with rhPCSK9_207-223_ VLP, a bivalent rhPCSK9 VLP vaccine, or, as a control, wild type Qß VLPs. Plasma was obtained after three immunizations (at day 77) and **b** anti-PCSK9_207-223_, **c** anti-PCSK9_153-163_, and **d** anti-full length hPCSK9 IgG levels were measured by ELISA. Data represents mean ELISA values for each group of macaques, error bars show the standard error of the mean.

After the third immunization, anti-PCSK9 antibody responses were measured by ELISA. Both rhPCSK9 VLP vaccines, but not the control vaccine, generated strong anti-PCSK9 IgG responses against rhPCSK9 peptides (Fig. 6b and 6c) and full-length hPCSK9 protein (Fig. 6d). Although anti-PCSK9 antibody titers declined over the course of the study, these titers remained high over four months following the initial immunization (Fig. S2).

### Immunization with PCSK9 VLPs decreases LDL-C levels in NHPs

To test vaccine efficacy, we measured lipid levels at multiple timepoints after immunization, focusing on LDL-C levels. LDL-C levels in NHPs immunized with control Qß VLPs were statistically similar to baseline levels at all timepoints tested, even after the macaques in this group were administered statins (Fig. 7). In contrast, we observed a statistically significant 28% decrease in plasma LDL-C levels in macaques that received the bivalent rhPCSK9 VLP after two immunizations (at day 42). LDL-C levels in this group were not affected by statin administration (which began on day 77) and remained below baseline levels until the day 133 timepoint (Fig. 7). Similar to what we observed previously in a pilot NHP study^30^, LDL-C levels in NHPs immunized with rhPCSK9_207-223_ VLPs alone did not decrease until after these macaques received statins (beginning on day 91; Fig. 7). In this group, LDL-C began rising once statins were withdrawn. Apolipoprotein B (ApoB), the major protein component of LDL that serves as the ligand for LDL-R, was also significantly reduced relative to controls in NHPs that received two immunizations (at day 42) with the bivalent rhPCSK9 VLP vaccine, but not in NHPs that were vaccinated with rhPCSK9_207-223_ VLPs (Fig. S3). After treatment with statins (at day 91), ApoB levels were reduced in both experimental groups relative to controls, but these differences were not statistically significant (Fig. S3). HDL-C levels did not change significantly within any of the groups over the course of the study (Fig. S3).

**Fig. 7.**
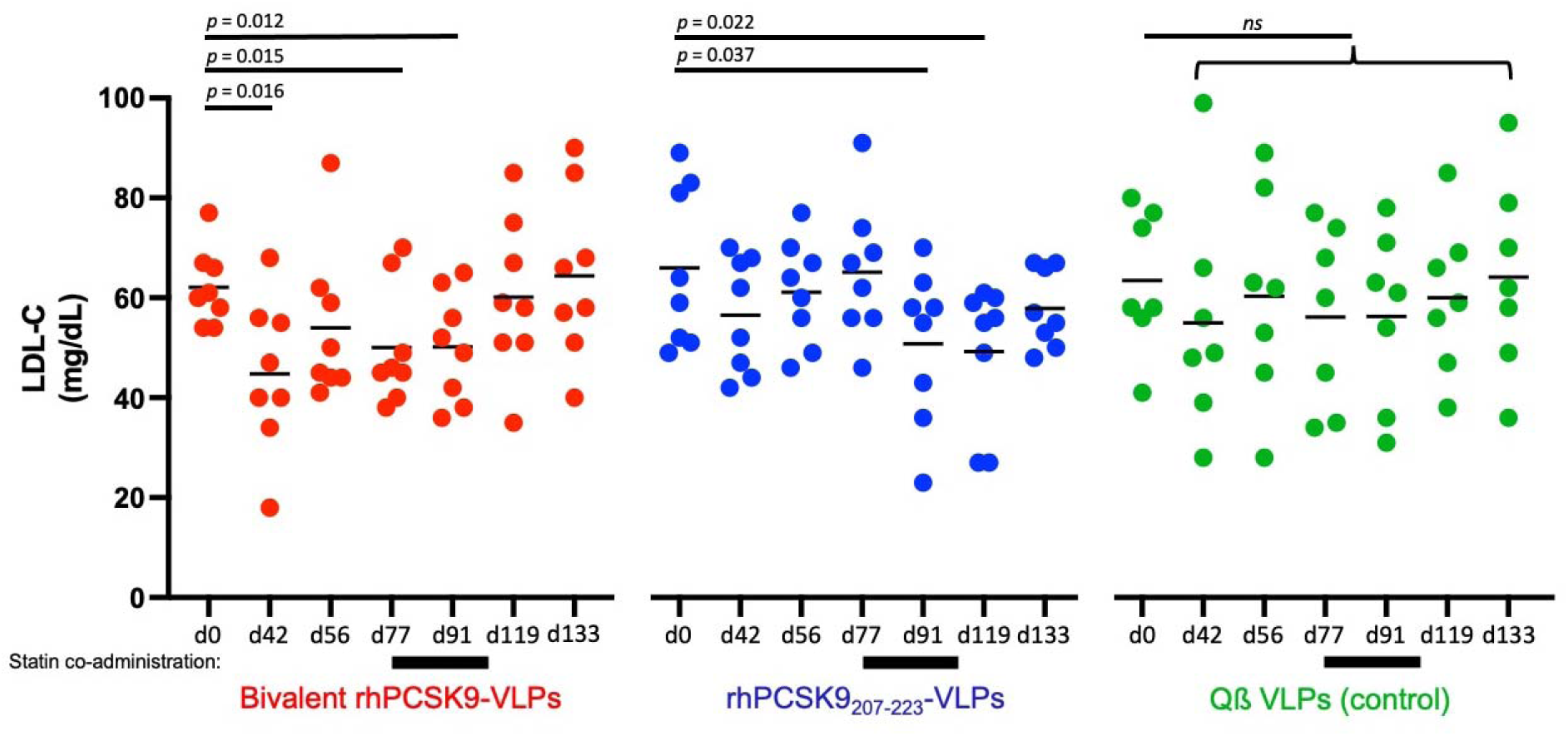
Plasma LDL-C levels in immunized rhesus macaques. Plasma LDL-C levels at different days post-prime in macaques immunized with bivalent rhPCSK9 VLPs (red), rhPCSK9_207-223_ VLPs (blue) or, as a control, wild type Qß VLPs (green). Macaques were treated daily with simvastatin from day 77-105 (represented by black bars below the x-axis). Each data point represents an individual animal, lines represent mean values for each group of macaques. Significance was determined by one-tailed t test.

### Single and bivalent PCSK9 vaccines differentially affect circulating plasma PCSK9 levels in immunized NHPs

We evaluated the effects of immunization on circulating plasma PCSK9 levels at day 77, after macaques had received three immunizations but prior to statin administration (which has been shown to increase circulating PCSK9^38–40^). As we observed in mice, plasma PCSK9 levels were elevated in the NHPs that were immunized with the single component vaccine, rhPCSK9_207-223_ VLPs (Fig. 8a). In contrast, NHPs that received the bivalent rhPCSK9 VLP vaccine had lower circulating PCSK9 levels, similar to the control group. Nearly identical plasma PCSK9 levels were measured at necropsy (Fig. 8b), after the NHPs had received an additional booster immunization. Thus, unlike in mice, the bivalent PCSK9 vaccine did not significantly reduce plasma PCSK9 levels relative to controls in macaques.

**Fig. 8.**
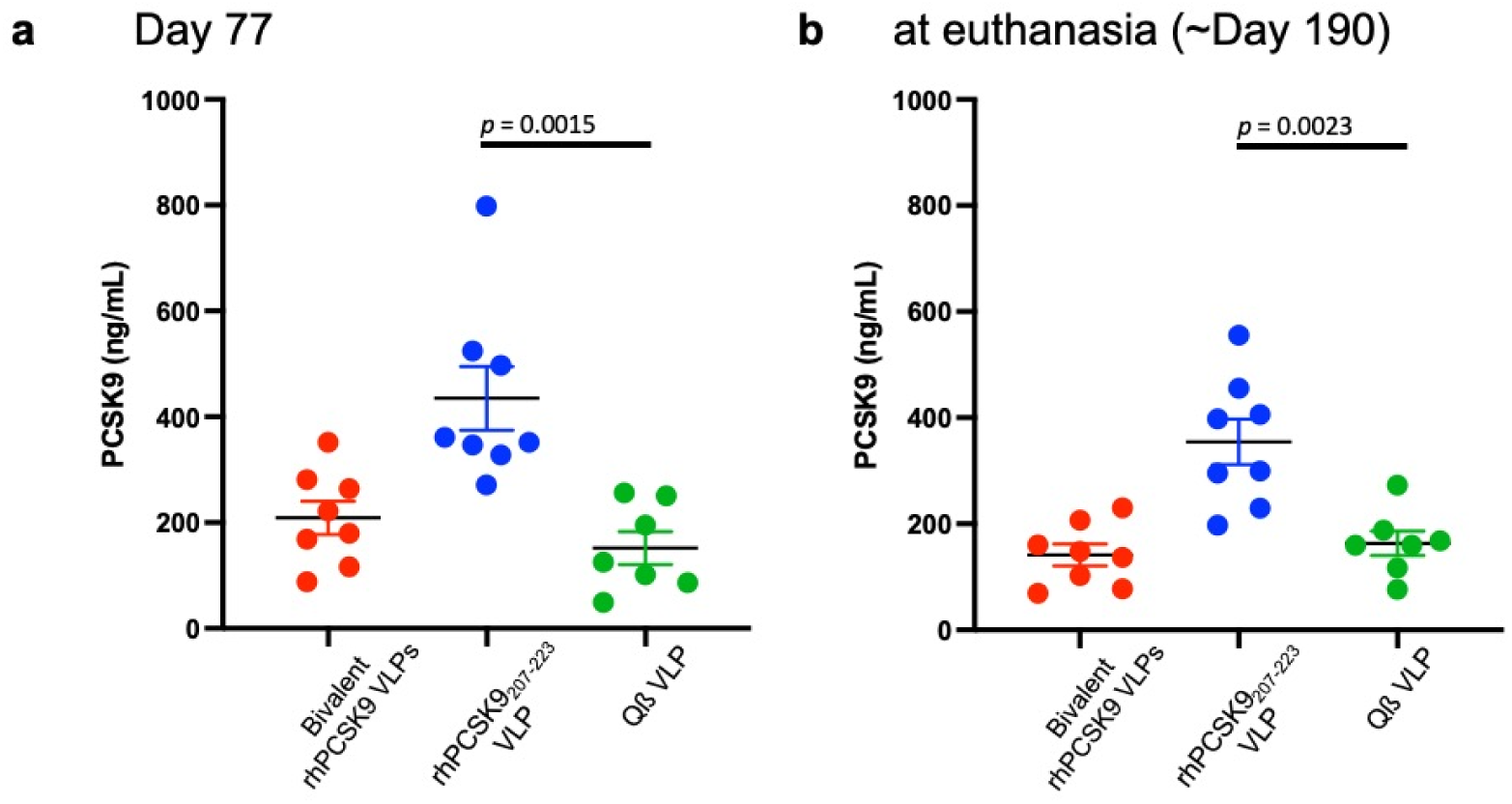
Circulating PCSK9 levels in immunized rhesus macaques. Plasma was collected three weeks following the third immunization (day 77, **a**) or following a fourth immunization at necropsy (day 190, **b**). Circulating PCSK9 levels were determined by ELISA. Each data point represents an individual animal, lines represent mean values for each group of macaques, and error bars show SEM. Significance was determined by two-tailed t test.

### Boosting increases anti-PCSK9 antibody levels, lowers LDL-C, and increases liver expression of LDL-R

At 161 days after the initial prime, and about a month prior to necropsy, NHPs received a booster immunization (Fig. 6a). Boosting increased anti-PCSK9 IgG titers by ∼5-to 10-fold in both groups which received rhPCSK9 VLP vaccines (Fig. 9a). We also evaluated the effects of boosting on plasma LDL-C by comparing lipid levels at the booster timepoint (day 161) with LDL-C levels in plasma taken 2- and 4-weeks later (at day 175 and at necropsy). As is shown in Fig. 9b, boosting significantly reduced LDL-C levels in macaques that received the bivalent rhPCSK9 VLP vaccine, but not in the groups that received the individual vaccine or controls.

**Fig. 9.**
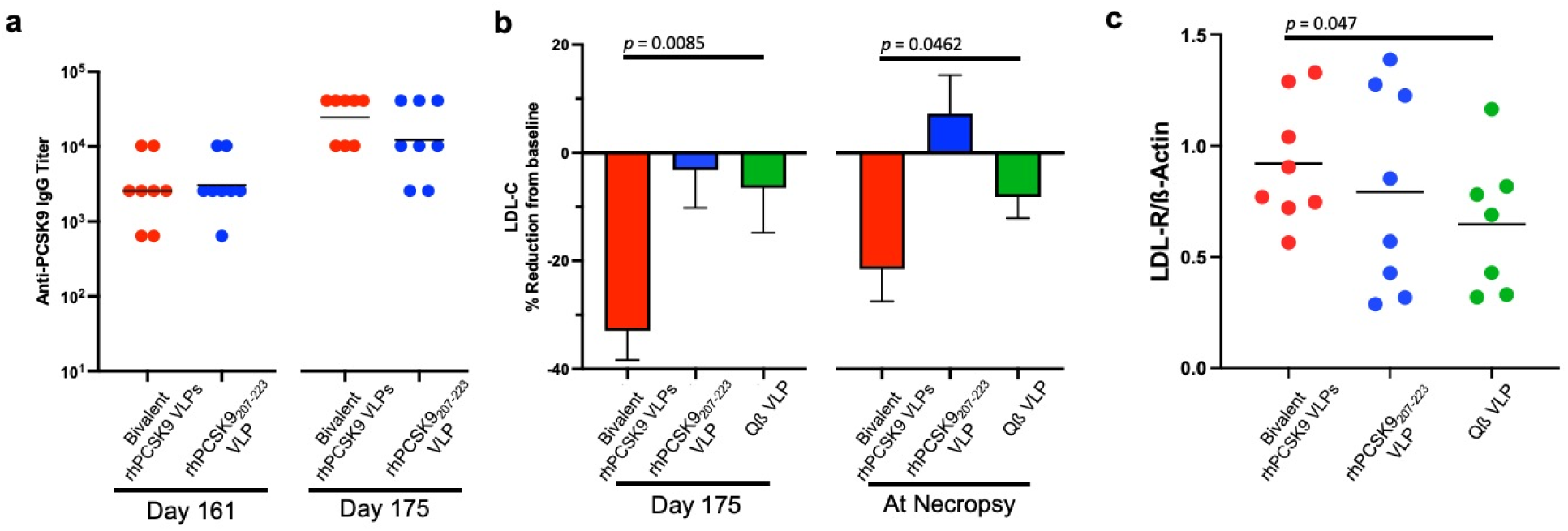
Boosting increases anti-PCSK9 antibody levels, lowers LDL-C, and increases expression of LDL-R in the liver. **a** Anti-PCSK9 IgG levels were measured by ELISA prior to boosting (at day 161) and two weeks following the boost (day 175). **b** Plasma LDL-C levels were measured at day 175 and at necropsy and were compared to baseline (at d161, prior to prime). **c** LDL-R protein levels were measured by Western blot (shown in Fig. S5), quantified using Image J, and then normalized by comparing the expression of LDL-R to β-actin (as a loading control). Each data point represents an individual macaque and means for each group are shown using a black line. Experimental groups were compared statistically by one-tailed t test.

At approximately day 190, macaques were euthanized and subjected to a full necropsy. We did not observe any gross pathology in any of the animals in this study. To further investigate the effects of anti-PCSK9 antibodies, we measured liver LDL-R expression. Cell lysates from liver samples were evaluated for LDL-R expression by Western blot analysis. As was previously observed in mice, there was a significant increase in liver LDL-R expression in the NHPs that were immunized with the bivalent rhPCSK9 VLP vaccine relative to the control group (Fig. 9c). NHPs that were immunized with rhPCSK9_207-223_ VLPs had slightly higher LDL-R levels, but this change was not statistically significant. To look at possible effects of vaccination on gene expression in the liver, mRNA levels of several genes involved in cholesterol metabolism, including *LDLR*, *APOB*, *HMGCR*, *HNF1A*, and *SREBF-2,* were measured. These mRNA levels were not significantly changed in vaccinated groups relative to controls (Fig. S4).

## Discussion

The ability to induce antibodies against self-antigens is seemingly limited by the mechanisms of immune tolerance. However, we and others have shown that multivalent display of antigens on the surface of nanoparticle-based platforms, such as VLPs, can efficiently induce antibodies against self-antigens involved in human disease, including PCSK9^27, 41^. Previously, we evaluated the immunogenicity and cholesterol-lowering efficacy of VLP-based vaccines displaying discrete peptide epitopes from human PCSK9^30^. These studies identified several PCSK9 epitopes that, when displayed on VLPs, elicited strong anti-PCSK9 IgG antibody responses and lowered total cholesterol levels in mice. The lead vaccine candidate from that study, which displays a peptide representing human PCSK9 amino acids 207-223 on Qß VLPs (hPCSK9_207-223_ VLPs), not only decreased total cholesterol levels in mice by approximately 30%, but also could be used in synergy with statins to lower LDL-C levels in a small pilot group of NHPs.

Here, we extended these preliminary studies in several different ways. First, we evaluated the immunogenicity of VLP-based PCSK9 vaccines that targeted species-specific PCSK9 sequences and confirmed that these vaccines could break immunological tolerance. Second, we showed that PCSK9 VLP vaccines could lower cholesterol levels in the LDLR^+/-^ mice, which exhibit elevated cholesterol levels. Third, we compared the efficacy of vaccines targeting individual PCSK9 epitopes and bivalent vaccines targeting multiple epitopes and showed that there are important differences in how these vaccines affect circulating PCSK9 levels. Fourth, we measured the longevity of anti-PCSK9 antibodies in both mice and macaques. Lastly, we evaluated the efficacy of lead PCSK9 vaccines in lowering LDL-C in larger groups of NHPs, both with and without statin co-administration. Taken together, these studies identified a PCSK9 vaccine regimen that induces long-lived anti-PCSK9 antibody responses and effectively lowers circulating LDL-C in primates without requiring co-administration of statins.

In this study, we focused on two linear PCSK9 epitopes, amino acids 153-163 and 207-223, which are located on the face of PCSK9 that is involved in LDL-R binding. Several groups, including our own, have demonstrated that vaccines targeting these epitopes can induce anti-PCSK9 antibodies and lower cholesterol levels in mice^30, 32, 33, 42, 43^. We were especially interested in evaluating a bivalent vaccine targeting both epitopes, hypothesizing that a bivalent vaccine targeting critical epitopes on PCSK9 could more effectively inhibit its function. In mice, we showed that the bivalent vaccine elicited similar anti-PCSK9 antibody titers as vaccines displaying individual mPCSK9 epitopes. In addition, the bivalent mPCSK9 vaccine increased LDL-R expression in the liver, and lowered mouse cholesterol levels to a similar extent as the vaccine that only targeted mPCSK9_207-223_. A major difference between the two approaches is that the vaccine targeting a single PCSK9 epitope increased circulating PCSK9 levels, whereas immunization with the bivalent mPCSK9 vaccine resulted in a ∼33% reduction in serum PCSK9 compared to controls. These data echo recent work by Goksøyr and colleagues, which showed that immunization with a VLP displaying full-length PCSK9 could elicit a broad, polyclonal anti-PCSK9 antibody response that lowered circulating PCSK9 levels, but immunization with VLPs displaying shorter peptides (amino acids 210-226 or 156-227) increased free PCSK9 levels^32^. Taken together with other evidence in the literature, these data indicate that vaccines which induce monoclonal-like (or oligoclonal) antibody responses against discrete PCSK9 epitopes lead to the formation of antibody-antigen complexes that are not efficiently cleared, extending PCSK9 half-life in serum. In contrast, polyclonal antibody responses that target multiple distinct epitopes on PCSK9 enhance its clearance, presumably by facilitating uptake of immune complexes by phagocytic cells (because PCSK9 is bound by multiple antibody molecules). In mice, however, vaccine-mediated clearance of PCSK9 was not essential for cholesterol lowering activity; individual and bivalent vaccines lowered cholesterol levels similarly. This suggests that when PCSK9 is bound to an antibody in the circulation it is not active in promoting the internalization of LDL-R, most likely because the antibody sterically blocks its interaction with the receptor.

We also evaluated the two different PCSK9 vaccines in rhesus macaques, which have a lipid profile that is more comparable to humans than mice^44^. One important limitation of our studies was that the mean baseline LDL-C level in macaques was low, ∼60mg/dL, which is normal for healthy animals. Low baseline LDL-C levels may have limited the magnitude of LDL-C lowering upon vaccination. We were particularly interested in determining whether immunization against PCSK9 could lower LDL-C as a monotherapy, or, as we showed previously in our pilot study^30^, whether co-administration of statins was required. Whereas immunization with rhPCSK9_207-223_ VLPs only lowered LDL-C levels in combination with statins, the bivalent rhPCSK9 VLP vaccine effectively lowered LDL-C levels (by ∼25-30%) without requiring statin co-administration. Macaques in this group also had higher levels of liver LDL-R expression and lower levels of circulating ApoB and PCSK9, consistent with the observed enhanced LDL-C lowering activity. It is unclear why the bivalent vaccine was more effective at lowering LDL-C in the absence of statins; both vaccines were highly immunogenic and elicited similar anti-PCSK9 antibody levels. This suggests that it may be possible to achieve clinically meaningful reductions in LDL-C by vaccination even without the use of statins. Because many patients do not tolerate or are not compliant with statin use^10^, this has important practical implications for this possible therapeutic approach.

We also measured the longevity of anti-PCSK9 antibody responses. In mice, we observed that anti-PCSK9 antibody levels slowly declined over time, with a half-life of approximately 20 weeks after the initial antibody decay. These half-life data are comparable what we had observed previously with VLP-based vaccines targeting self-antigens in animal models^45, 46^, as well as data from human clinical trials of VLP-based vaccines targeting the self-antigens amyloid-beta^29^ and angiotensin II^28^. In NHPs, anti-PCSK9 antibody levels and LDL-C lowering effects also declined over time, but boosting increased anti-PCSK9 antibody titers and lowered LDL-C. Follow-up studies that correlate anti-PCSK9 antibody levels with LDL-C lowering will be critical for determining an optimal boosting schedule for PCSK9 vaccines.

The current clinically approved PCSK9 inhibitors are remarkably effective at lowering LDL-C levels. Nevertheless, there are major barriers that limit access to PCSK9 inhibitors, including their prohibitive expense and formulatory restrictions imposed by insurance providers^47^. Efforts to develop next generation PCSK9 inhibitors have focused on achieving increased reduction of LDL-C, enhancing the duration of effect, altering the mode of administration, and decreasing cost^15^. Promising emerging approaches include an orally delivered antisense oligonucleotide (ASO)^48^, synthetic adnectin polypeptides^49^, cyclic peptides that can be delivered orally^50, 51^, and an *in vivo* CRISPR-mediated base editing approach to permanently introduce loss-of-function mutations in PCSK9^52^. A vaccine-based approach may fill a niche as a long-lasting, inexpensive strategy for targeting PCSK9. Indeed, several different groups have now reported promising pre-clinical results from studies of PCSK9 peptide-based vaccines^32, 33, 42, 53–56^, although PCSK9 vaccines are generally less potent at lowering LDL-C than other therapeutic approaches. Two PCSK9 peptide targeted vaccines, AT04A and AT06A, have been evaluated in clinical trials. Three doses of AT04A, which is a 10-amino acid PCSK9 peptide linked to the carrier protein keyhole limpet hemocyanin (KLH), elicited anti-PCSK9 antibodies (albeit low titer responses: ∼1:134) and, at peak efficacy, resulted in an 11-13% reduction in LDL-C levels^57^. Even though these reductions were small, they would still be predicted to reduce the incidence of cardiovascular events^58^. The use of a more immunogenic vaccine platform than KLH would likely result in stronger antibody responses and greater reduction in LDL-C levels, as has been shown previously^59^.

In conclusion, we have shown that a bivalent vaccine consisting of VLPs displaying two different PCSK9-derived peptides can induce robust anti-PCSK9 antibody responses, decrease serum PCSK9 levels, increase liver expressed LDL-R, and efficiently lower cholesterol levels in mice and macaques. These findings strongly support the development of an alternative vaccine-based approach for inhibiting PCSK9 activity and lowering LDL-C.

## Materials and methods

### VLP production

Qβ bacteriophage VLPs were produced by transforming C41 *Escherichia coli* (*E. coli*) cells with a pET-derived plasmid encoding the Qß bacteriophage viral coat proteins. Transformed C41 cells were grown at 37°C using Luria Bertani broth containing 60 µg/mL kanamycin until the cells reached an OD600 of 0.6. Coat protein expression was induced using 0.4 mM isopropyl-β-D-1-thiogalactopyranoside (IPTG) and grown at 37 °C for 3 h. Cell pellets were collected and re-suspended in lysis buffer (50mM Tris-HCL, 100mM NaCl, 10mM EDTA, pH 8.5). Cells were lysed by sonication and cell lysates were clarified by centrifugation (15,000×*g*, 20min, 4°C). Soluble VLPs were purified by precipitation using 70% saturated (NH_4_)_2_SO_4_, followed by size exclusion chromatography (SEC) using a Sepharose CL-4B column. The column was pre-equilibrated with a purification buffer (40 mM Tris-HCl, 400 mM NaCl, 8.2 mM MgSO_4_, pH 7.4). VLPs were concentrated from SEC purified fractions by precipitation using 70% saturated (NH_4_)_2_SO_4_ and then extensively dialyzed versus PBS, pH 7.4.

### Display of PCSK9-derived peptides on VLPs

Peptides representing mPCSK9 sequences mPCSK9_153-163_ (SIPWNLERIIP) and mPCSK9_207-223_ (SVPEEDGTRFHRQASKC) and rhPCSK9 sequences rhPCSK9_153-163_ (SIPWNLERITP) and rhPCSK9_207-223_ (SVPEEDGTRFHRQASKC; the same as the mPCSK9_207-223_ sequence) were synthesized by GenScript. PCSK9_153-163_ peptides were modified with a C-terminal cysteine residue preceded by a 3-glycine-spacer sequence (-GGGC) to facilitate conjugation to VLPs. PCSK9_207-223_ peptides contain a naturally occurring C-terminal cysteine residue, so this peptide was not modified with a linker sequence. PCSK9 peptides were conjugated to Qß bacteriophage VLPs using succinimidyl 6-((ß-maleimidopropionamido)hexanoate) (SMPH; Thermo Fisher Scientific), as described previously^60^. Conjugation efficiency was measured using sodium dodecyl sulfate-polyacrylamide gel electrophoresis (SDS-PAGE).

### Animals and Immunizations

Studies using mice were performed in accordance with guidelines of the University of New Mexico Animal Care and Use Committees (protocol 19-200870-HSC). Immunizations were performed using 4-week-old Balb/c mice or LDLR^+/-^ mice, which were generated by breeding male LDLR^-/-^ mice (B6.129S7-*Lflr^tm1Her^*/J; The Jackson Laboratory) with female C57BL/6 mice and using first-generation heterologous offspring. Mice were immunized intramuscularly with 5µg of mPCSK9 VLPs without exogenous adjuvant. Mice were boosted twice at 3 and 6 weeks after the initial prime. Some groups of mice received a bivalent vaccine which consisted of 5µg of mPCSK9_153-163_ VLPs and 5µg of mPCSK9_207-223_ VLPs. Blood plasma was collected prior to the first immunization, two weeks following each immunization, and then three weeks following the final immunization. In some cases, plasma was also collected monthly for one year following the initial immunization. To minimize fluctuations in cholesterol levels, mice were fasted for 3 hours prior to each bleed. Immunization experiments were repeated several times and the data reported in this manuscript represent aggregated data from multiple experiments.

The non-human primate (NHP) studies used adult rhesus macaques (Macaca mulatta), between 7-12 years of age, from the breeding center of the California National Primate Research Center, which is negative for simian immunodeficiency virus, type D retrovirus and simian T-cell lymphotropic virus type 1. The CNPRC is accredited by the Association for Assessment and Accreditation of Laboratory Animal Care International (AAALAC). Animal care was performed in compliance with the 2011 *Guide for the Care and Use of Laboratory Animals* provided by the Institute for Laboratory Animal Research. The study was approved by the Institutional Animal Care and Use Committee of the University of California, Davis (study protocol 21066). For sample collections and immunizations, animals were sedated with ketamine anesthesia (10mg/kg body weight) administered by the intramuscular (IM) route.

Twenty-three NHPs were divided into three experimental groups, immunized with 1) 50µg of rhPCSK9_207-223_ VLPs (n=8), 2) a bivalent vaccine consisting of 25µg of each rhPCSK9_153-163_ VLPs and rhPCSK9_207-223_ VLPs (n=8), and 3) 50 µg wild-type Qß VLPs (n=7). NHPs were immunized intramuscularly with vaccine formulated with 2% Alhydrogel hydroxide at a 1:1 (volume:volume) ratio on days 0, 28, and 56. Blood plasma was collected two weeks following each immunization. Beginning on day 77, NHPs were given oral simvastatin daily (at 30 mg/kg/day) through day 105. Based on each individual animal’s weight, a full number of 80 mg simvastatin tables, calculated to approximate the 30 mg/kg dose, were crushed to powder, that was mixed with food supplements that were provided to the animals; uptake was monitored and recorded. On days when animals were sedated for procedures, the simvastatin was administered via orogastric intubation. The number of tablets was adjusted weekly based on most recent body weights. NHPs received another dose of vaccine at day 161, approximately 30 days prior to euthanasia (via an overdose of pentobarbital). A necropsy was performed to evaluate gross pathology and collect tissues.

### Measuring antibody titers

PCSK9-specific IgG titers were determined by end-point dilution ELISA, using either PCSK9 peptides or recombinant mPCSK9/hPCSK9 protein (R&D Systems) as the coating antigen. Because rhPCSK9 is not commercially available, hPCSK9 was used as the target antigen for evaluating antibody responses in macaques immunized with rhPCSK9 epitopes. For peptide ELISAs, Immulon 2 plates (Thermo Fisher Scientific) were coated with 500ng streptavidin (Invitrogen) for 2 hours at 37°C. Following washing, SMPH was added to wells at 1 μg/well and incubated for 1 hour at room temperature. Specific peptides were added to the wells at 1 μg/well and incubated overnight at 4°C. For full-length PCSK9 ELISAs, plates were coated with 250ng of PCSK9 and incubated at 4°C overnight. For all ELISAs, wells were blocked with PBS-0.5% nonfat dry milk for 2 hours at room temperature. Plasma isolated from immunized animals were serially diluted in PBS-0.5% milk, applied to wells, and incubated at room temperature for 2.5 hours. Reactivity to the target antigen was detected using HRP-labeled goat anti-mouse IgG (Jackson Immunoresearch, diluted 1:4000) or goat anti-monkey IgG (Fitzgerald Industries International Inc; diluted 1:4000) for 1 hour. The reaction was developed using TMB substrate (Thermo Fisher Scientific) and stopped using 1% HCl. The optical density was measured at 450 nm using a microwell plate reader (accuSkan FC; Fisher Scientific). End-point dilution titer was defined as the greatest sera dilution that yielded an OD_450_ value >2-fold over background.

### Plasma lipid quantification

Plasma lipids were measured enzymatically using a ChemWell instrument and Roche reagents.

### Plasma PCSK9 quantification

Plasma PCSK9 levels were measured using mouse or human PCSK9 ELISA kits (R&D Systems) by comparing experimental samples diluted 1:20 (macaque samples) or 1:200 (mouse samples) to an internal standard curve.

### Measuring LDL-R expression in the liver

Mouse or rhesus macaque liver samples (∼100mg) were homogenized using the RIPA Lysis Buffer System (Santa Cruz Biotechnology) following the manufacturer’s protocol. Liver lysates were collected following centrifugation at 10,000x*g* for 30 minutes. Protein samples were separated by SDS-PAGE and transferred onto nitrocellulose membranes (Bio-Rad). Membranes were probed using goat anti-mouse LDLR at a 1:500 dilution (AF2255; R&D Systems) or goat anti-human LDLR at a 1:500 dilution (AF2148; R&D Systems) followed by incubation with an anti-goat IgG horseradish peroxidase-conjugated secondary antibody at a 1:1000 dilution (HAF107; R&D Systems). As a control, blots were also probed with an HRP-labelled rabbit anti-ß-actin monoclonal antibody at a 1:1000 dilution (#5152; Cell Signaling Technology). Western blots were visualized with SuperSignal West Pico Chemiluminescent reagents (Thermo Fisher Scientific) and band intensities were quantified using Image J software.

### Quantifying liver transcript levels

Transcripts were measured by quantitative (q) RT-PCR. Rhesus macaque liver samples (∼100mg) were homogenized using the lysis buffer provided with the PureLink RNA Mini Kit (Life Technologies) using the FastPrep-24 kit (MP Bio). RNA was amplified using the TaqPath 1-Step RT-qPCR Master Mix with gene-specific primer/probe sets purchased from Life Technologies. RNA levels of specific genes were measured by quantitative PCR using QuantStudio 7 (Thermo Scientific).

### Statistics

All statistical analyses of data were performed using GraphPad Prism 9. The specific statistical tests used are listed in the Figure legends.

### Data availability

The data sets used and/or analyzed in the current study are available from the corresponding author upon reasonable request.

## Acknowledgements

This research was funded by NIH grant R01HL131696, by the Intramural Research Program of NHLBI, and by an award from the National Center for Advancing Translational Sciences, National Institutes of Health under grant number UL1TR001449, and award P51OD011107 from the Office of Research Infrastructure Program, Office of The Director, National Institutes of Health to the California National Primate Research Center. We thank the staff of CNPRC Colony Management and Research Services, Clinical Laboratories, Anatomic and Clinical Pathology and Veterinary staff for their expert technical assistance. We also thank Alexandra Francian for her valuable comments on the manuscript.

## Supplement

**Supplementary Table 1.**
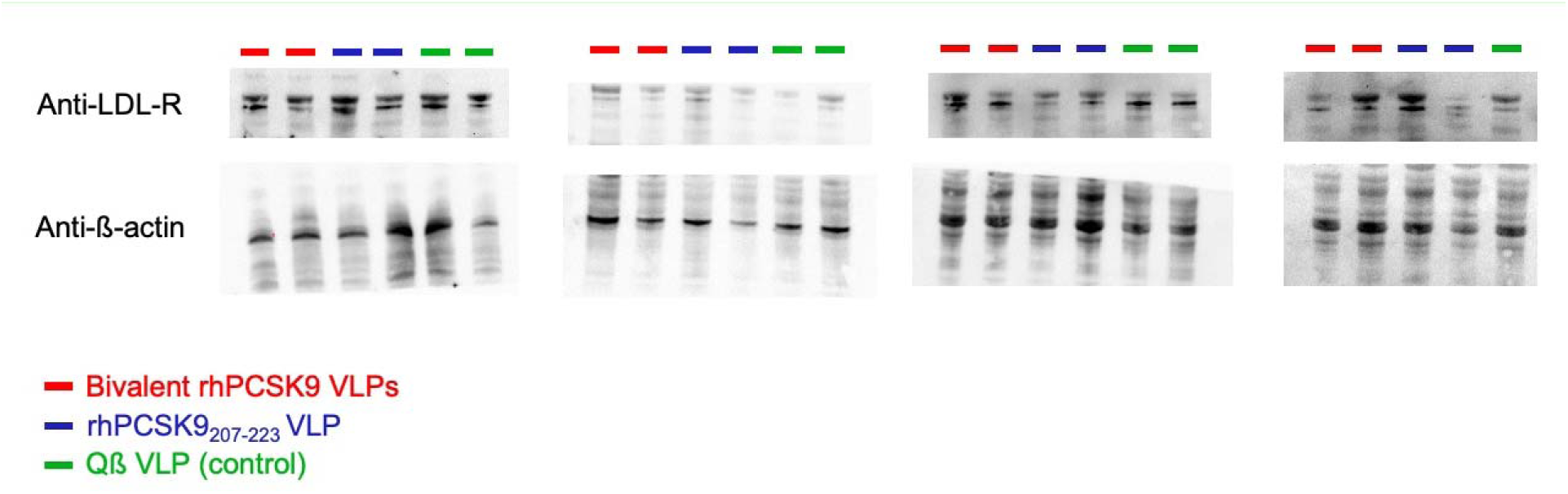
Baseline characteristics of NHPs enrolled in this study^1^

**Fig. S1.**
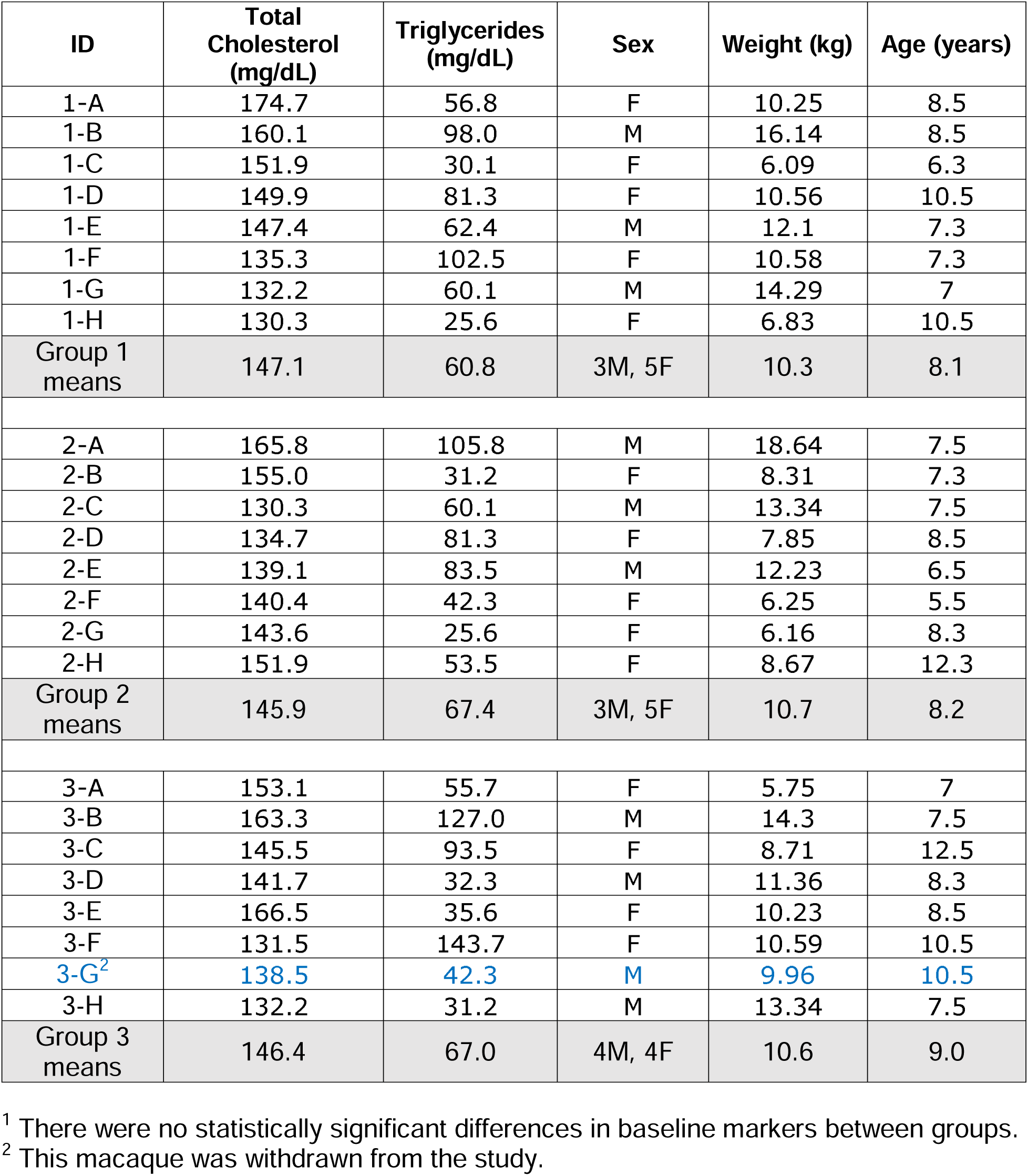
Western blot data graphed in. Fig. 4. Liver expression of LDL-R and ß-actin were measured by Western blot, samples obtained from mice immunized with Qß VLPs (lanes denoted with green bars) or bivalent mPCSK9-VLPs (red bars)

**Fig. S2.**
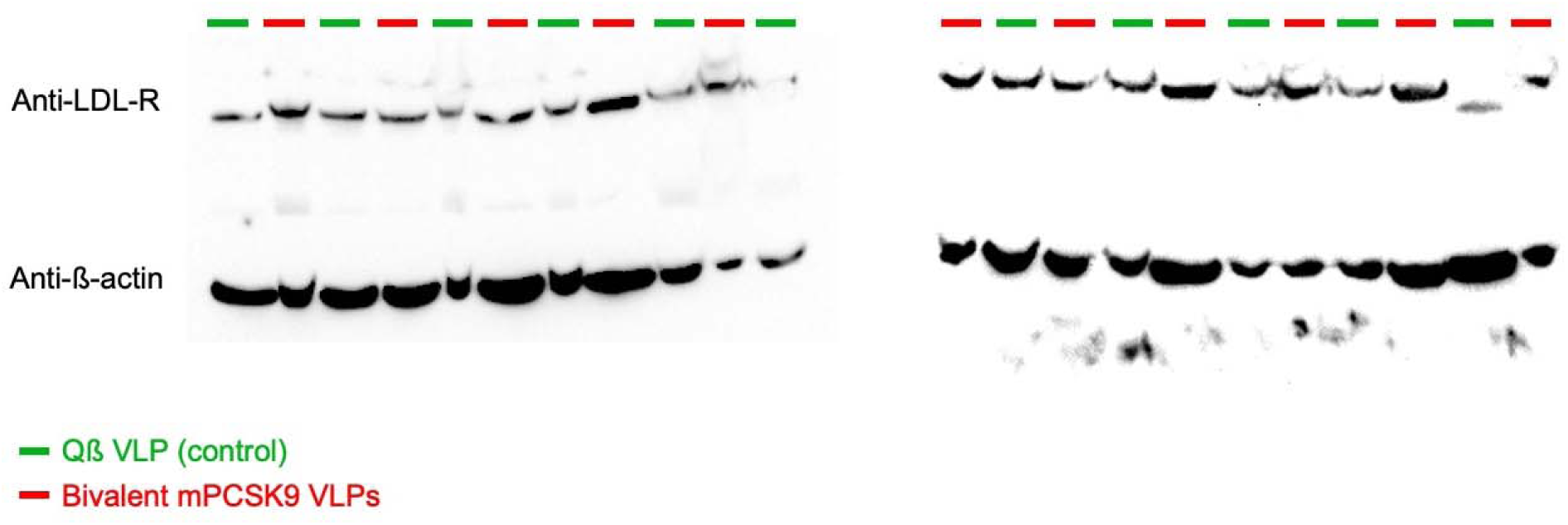
Longevity of anti-PCSK9 antibody responses in immunized rhesus macaques. Plasma was obtained at day 91 and day 133 of the study. Endpoint dilution IgG titers against full-length hPCSK9 were measured by ELISA. Each data point represents an individual macaque and the geometric mean titer for each group is shown using a black line.

**Fig. S3.**
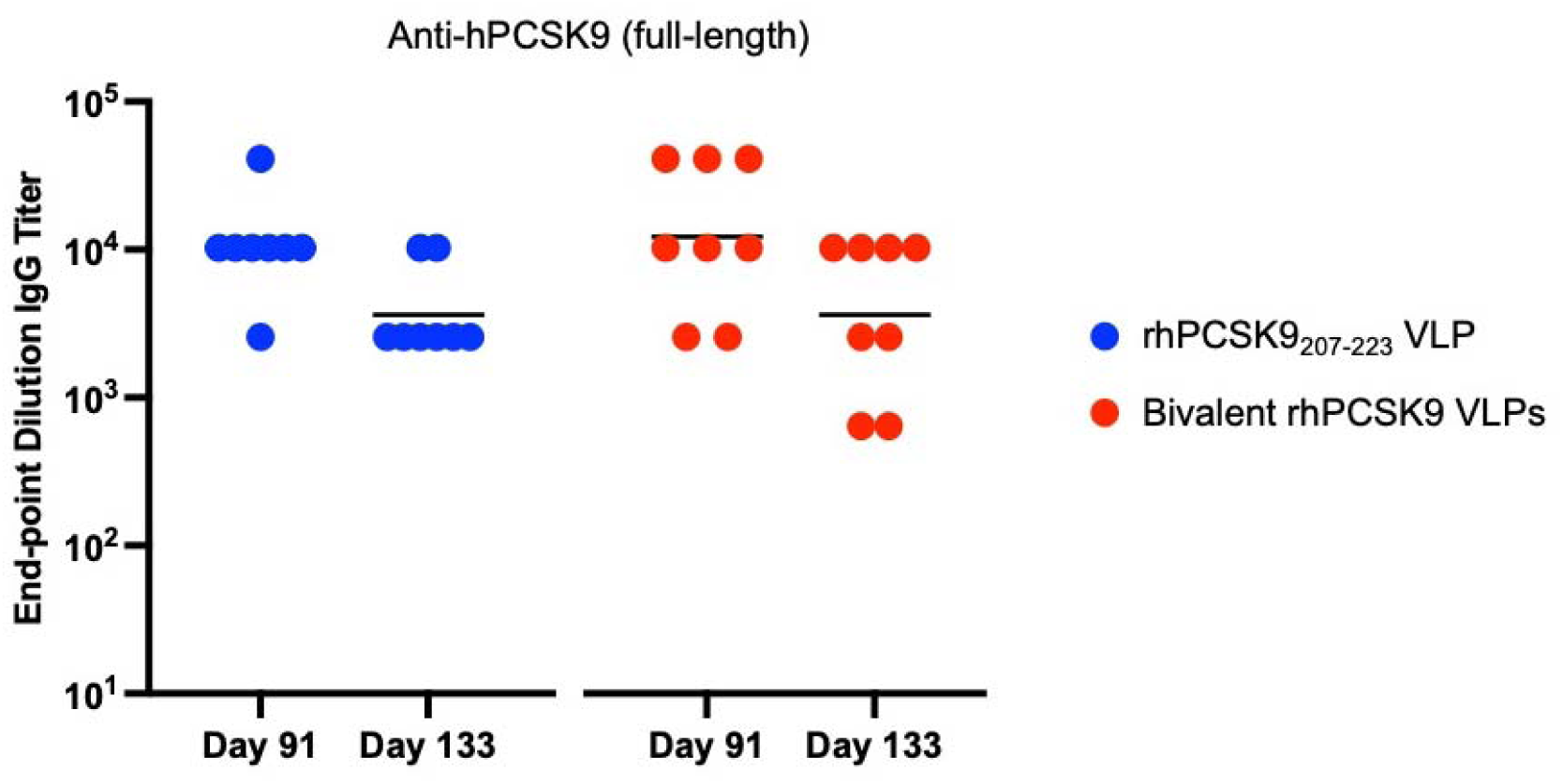
Plasma ApoB and HDL-C levels in immunized rhesus macaques. Plasma was obtained after two immunizations (day 42, **a** and **b**) and two weeks after daily simvastatin administration began (day 91, **c** and **d**). Plasma ApoB (**a** and **c**) and HDL-C (**b** and **d**) levels were measured and compared to baseline (at d0, prior to prime). Bars represent mean % reduction from baseline, error bars show SEM. Significance was determined by one-tailed t test.

**Fig. S4.**
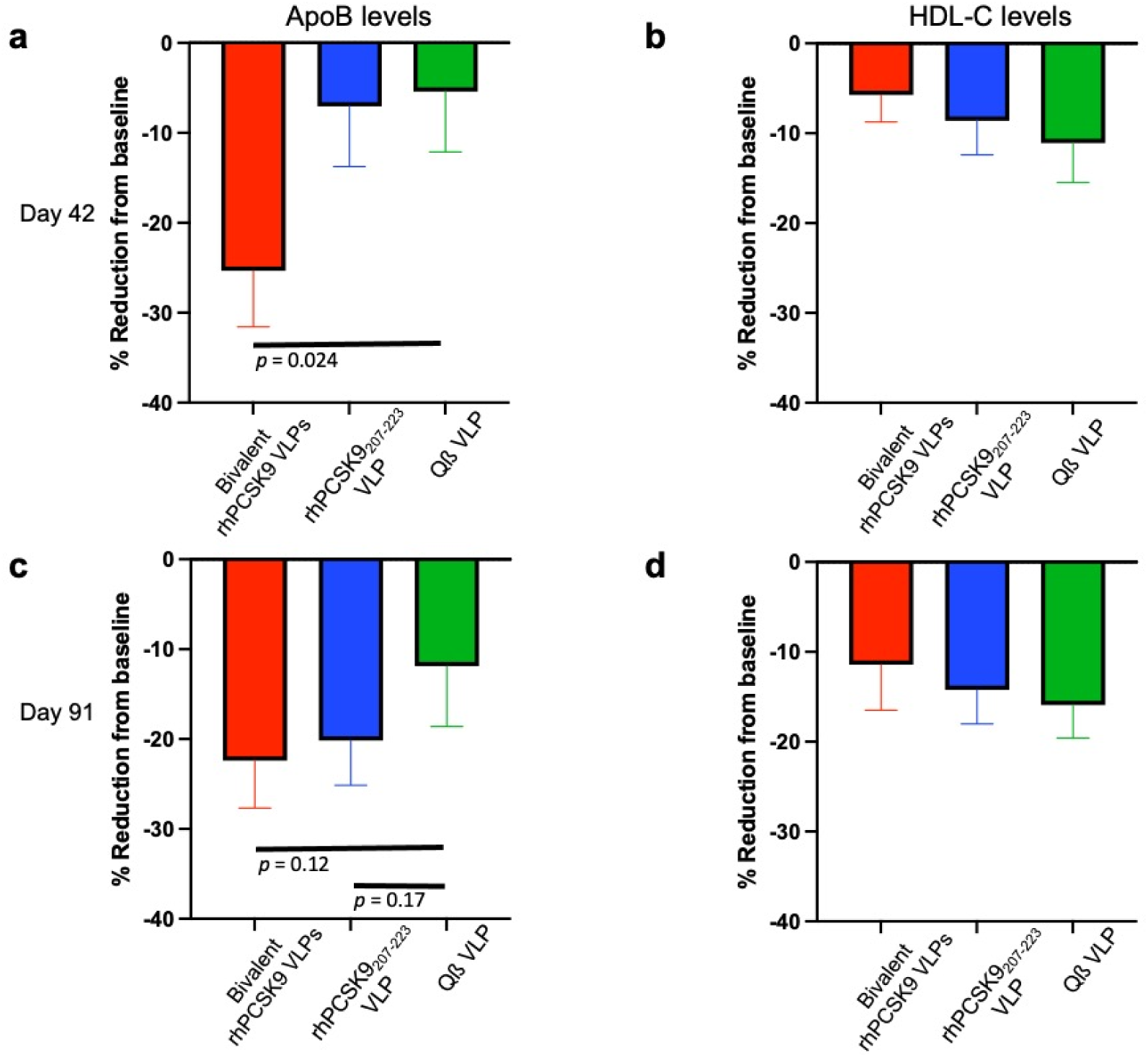
Relative mRNA expression in vaccinated groups of rhesus macaques. Liver mRNA levels were determined by qRT-PCR. Cycle threshold (Ct) values were determined for each mRNA species, normalized using a housekeeping mRNA (Actin) to determine a delta CT (dCT) value, and then compared to the mean control dCT value from the group immunized with Qß VLPs, to determine the fold change of mRNA expression.

**Fig. S5.**
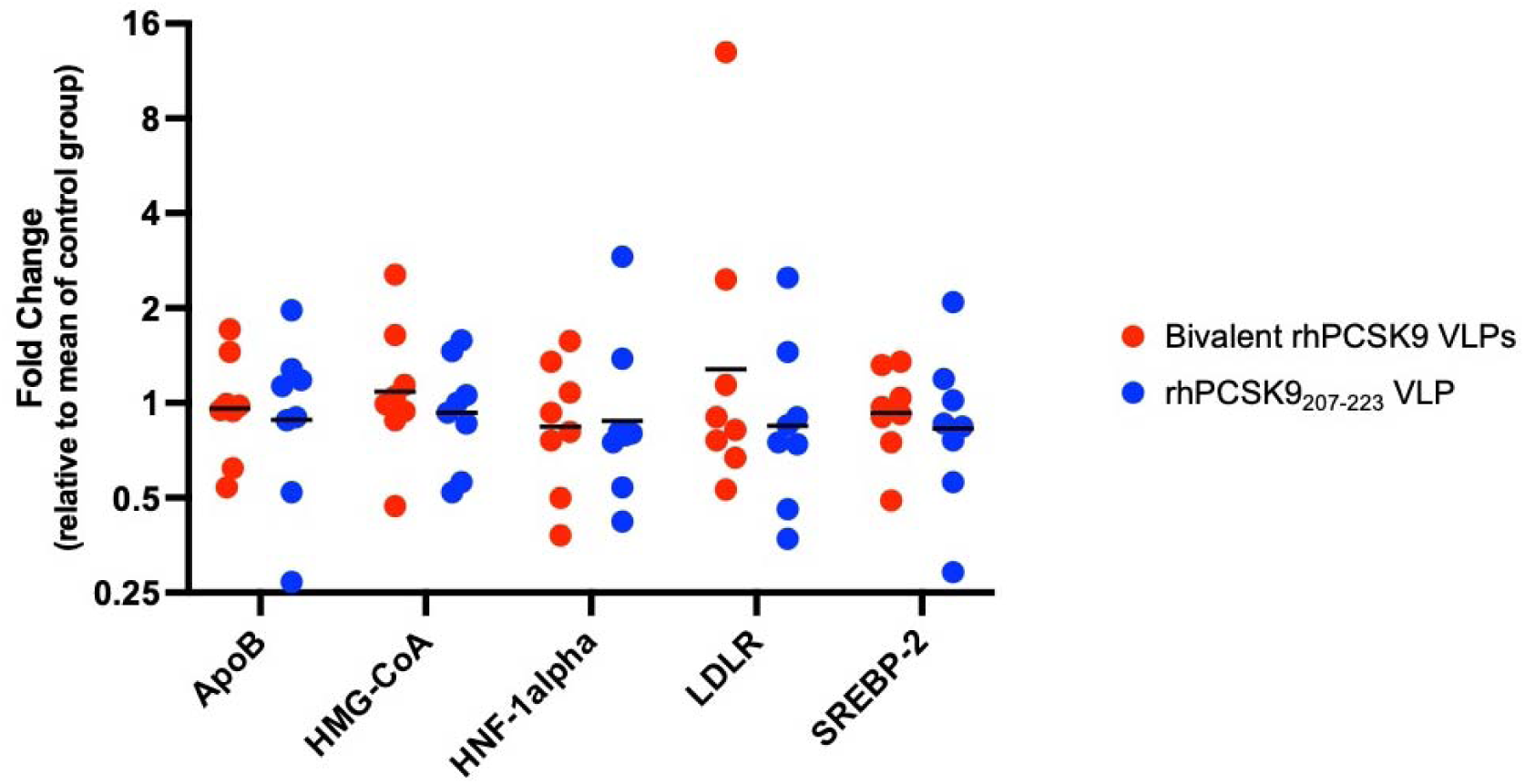
Western blot data graphed in. Fig. 9d. Liver expression of LDL-R and ß-actin were measured by Western blot, samples obtained from rhesus macaques immunized with bivalent rhPCSK9-VLPs (lanes denoted with red bars), rhPCSK9_207-223_-VLPs (blue bars), or Qß VLPs (green bars).

